# Rapid thermoforming of polycarbonate cell culture accessories from 3D printed molds

**DOI:** 10.1101/2025.07.07.663502

**Authors:** Ganesh Malayath, Nathaniel Huebsch

## Abstract

Bespoke cell culture devices are essential for tissue engineering applications. Traditional manufacturing methods for cell culture accessories involve injection molding and machining, which are too costly and time-consuming to implement for producing custom designs in small batches, and/or while testing the usefulness of a new design before mass producing it. Materials typically used for rapid design iteration, like poly(dimethylsiloxane) (PDMS) elastomers, surmount this issue but present new challenges of affinity for hydrophobic small molecules and sub-optimal interactions with sensitive cell types.

Here, we propose polycarbonate (PC) thermoforming as a solution for creating customized transparent and autoclavable accessories. We demonstrate that optimized preheating of PC overcomes issues with bubbling during thermoforming. The use of high heat deflection temperature (HDT) resins allows these PC devices to be thermoformed off molds created by Digital Light Processing (DLP) 3D prints, enabling rapid prototyping of the PC. Using this approach, we fabricated custom PC well plate inserts. These inserts combine many advantages of tissue culture polystyrene (negligible absorption of hydrophobic molecules, transparency, rigidity) and elastomers (ease of creating bespoke devices, ability to be sterilized by autoclaving) and are compatible with a variety of cell biology applications, including human induced pluripotent stem cells (iPSC) culture. PC inserts also supported iPSC differentiation into cardiomyocytes (iPS-CM) and micro-patterning of iPS-CM into cardiospheres. This low cost, customizable approach holds promise for a variety of bioengineering applications.

## INTRODUCTION

Discrepancies between the way cells respond to chemical cues (e.g. drugs) and genetic mutations in cell culture versus in the body have led the biomedical science community to push toward the development of “*in vivo-*like” cell culture platforms^1–6^ in which cells are subjected to biophysical stimuli that mimic the stimuli they would encounter in the body. For example, in the context of cardiovascular tissue engineering, neonatal rodent cardiomyocytes or human induced pluripotent stem cell (iPSC) derived cardiomyocytes (iPS-CM) are formed into organized, aligned tissues, or cardiospheres using various macro- and micro-scale devices (**Fig. 1**)^7–11^. These tissues may be further coaxed toward a more mature state via continuous mechanical^12^ or electrical stimulation^13,14^ pacing. The need to provide these geometric and/or biophysical cues is challenging within conventional cell culture platforms, often creating a need to create bespoke devices.

**Fig. 1.**
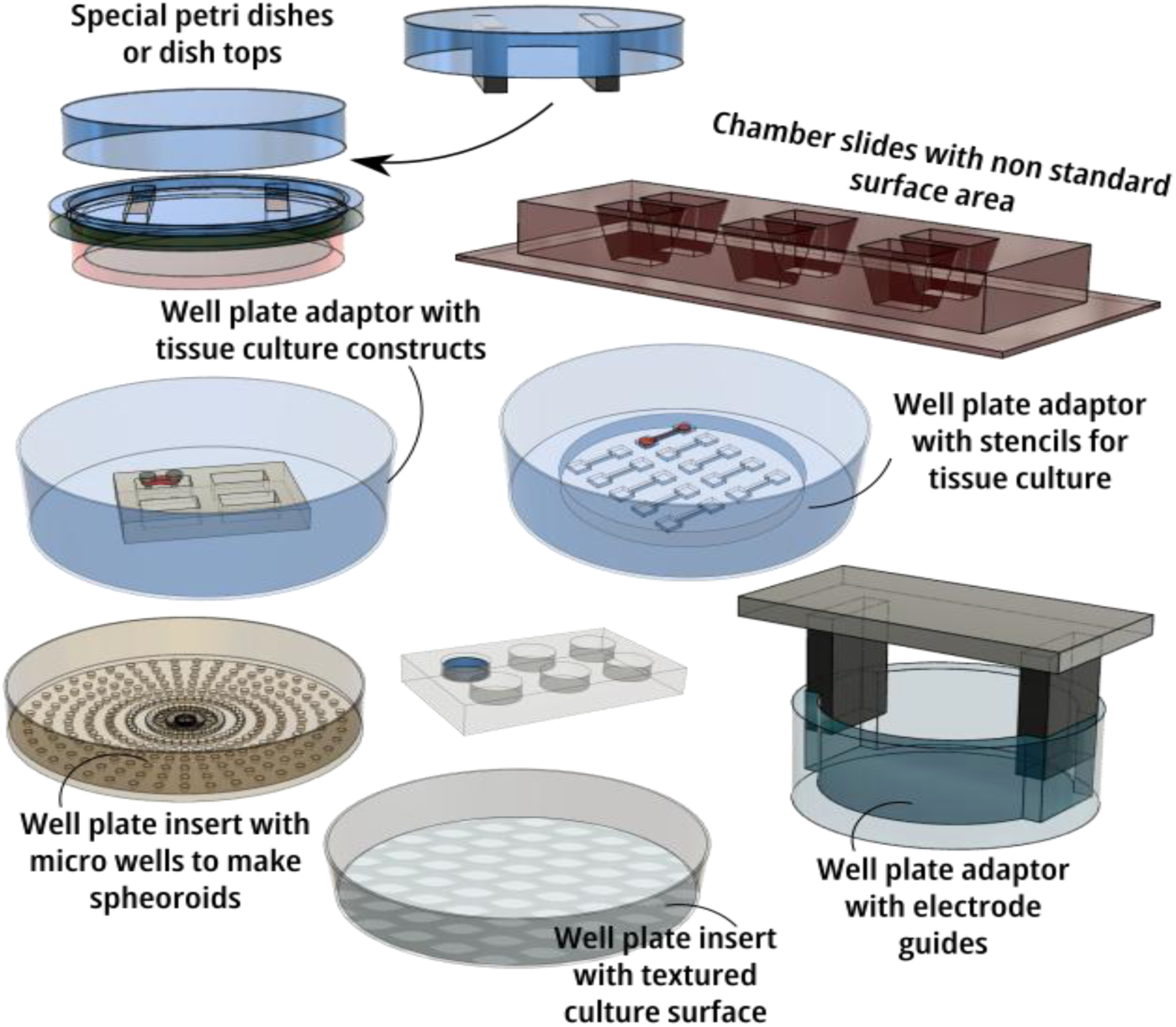
Specialized cell culture accessories are essential tools for bioengineering research: Schematic representation of customized cell culture/ tissue engineering systems used for maturing tissues with mechanical/electrical stimulation, studying tissue mechanics and live imaging with high resolution microscopy. *Schematics created using Inkscape vector graphics editor and Autodesk Fusion360*.

Conventional cell culture platforms, such as well plates, culture flasks, and chamber slides, are predominantly mass-produced using polystyrene (PS) through injection molding techniques and precision-machined tooling^15^. These standardized products, while widely available and highly efficient in supporting cell growth, offer limited customization options. Moreover, as polystyrene does not withstand autoclaving temperature, it is also challenging to attach customized devices or tissue culture constructs directly to these platforms without compromising sterility, especially for long-term studies.

To address the limitations of these commercial platforms, researchers commonly employ poly(dimethylsiloxane) (PDMS) to create bespoke inserts, adaptors, or molds to form engineered tissues (**Fig. 1**). These custom devices are made by crosslinking the PDMS elastomer over molds created through machining, 3D printing ^16^or photolithography ^17^. However, this approach presents its own challenges: 1) To get optical clarity necessary for imaging, the mold has to be polished extensively^18^; 2) PDMS exhibits an affinity for hydrophobic molecules, which can complicate drug screening efforts^19^; 3) for compatibility with high resolution microscopy, which demands very small working distances, PDMS devices must be thin, which introduces challenges in handling and device production^20^. Once designs are finalized, creating permanent molds for PDMS casting is challenging; conventional machining is costly and time-consuming^21^, while 3D printing, though more flexible and accessible, can lead to acrylate monomer leaching from resins into PDMS, inhibiting curing^22^. Soft lithography is useful in creating micro patterns and channels but can be challenging to use for making large accessories^23^. These limitations underscore the need for alternative materials and fabrication methods that can provide rapid, cost-effective, and biocompatible solutions for customized cell culture accessories.

Thermoforming is a versatile manufacturing process widely used in packaging, automotive, and dental applications^24^. In thermoforming, plastic sheets are heated until pliable, then formed over a mold using vacuum or pressure **(Fig. 2)**. The availability of low-cost (<$100) commercial small-scale thermoforming machines, popularized by dental guard production^25^, makes this technology widely accessible. Compared to methods that require elastomer crosslinking (e.g. traditional soft lithography), the rapid nature of thermoforming allows users to generate multiple parts within a short time (one minute/part) via repetitive use of the same mold.

**Fig. 2.**
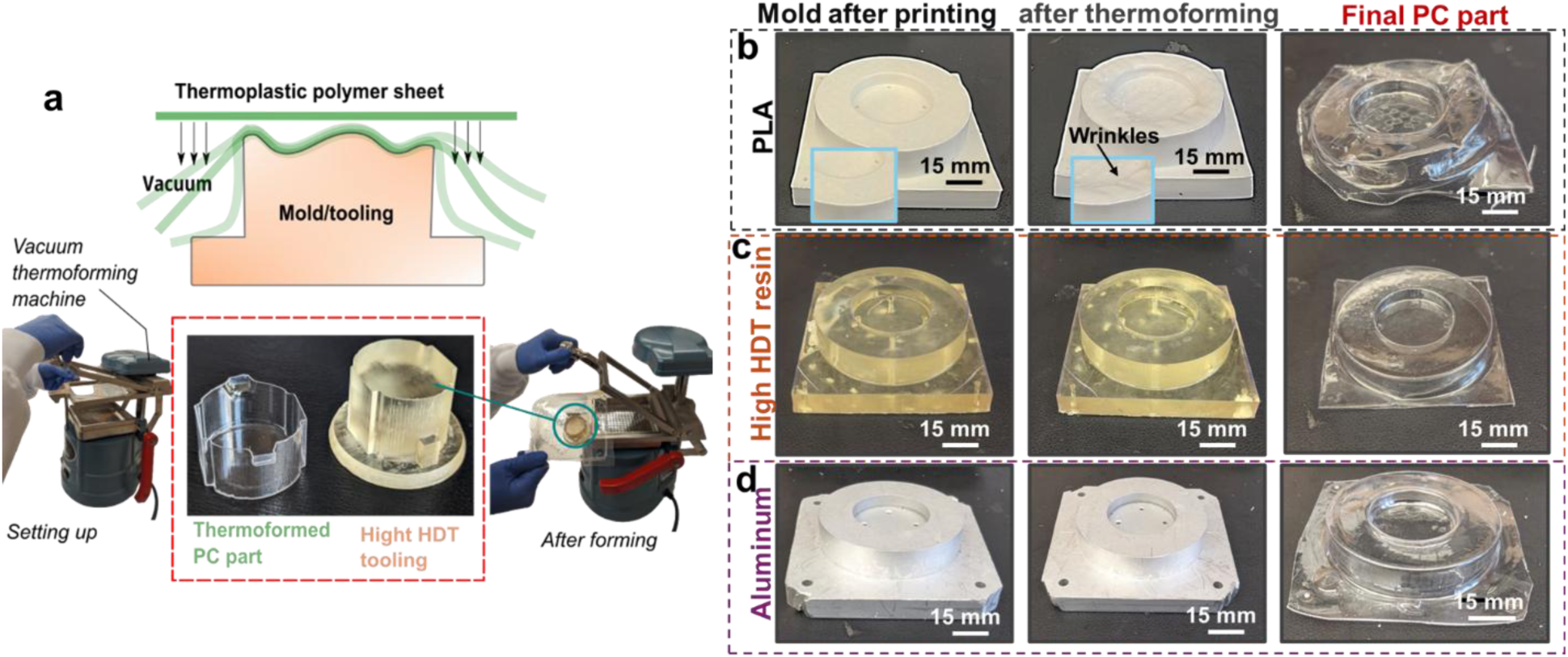
PC Thermoforming off Molds 3D-Printed from HDT Resins. **A)** Principle of thermoforming and proposed tooling method **B-D)** performance of different mold materials (B: poly-lactide acid, PLA; C: HDT Resin; D: 6061 aluminum) in thermoforming customized 70 mm petri dish top. Insets in panel B denote damage to the PLA mold after thermoforming.

While thermoforming is promising, adapting this technique to create bespoke cell culture accessories presents multiple challenges, which have limited application of this technique to micro-device fabrication^26^. First, cell culture accessories must be sterilized (preferably by autoclaving) and support cell culture (either as a substrate or indirectly as a housing). Sterilization by autoclaving, which is more widely available than gamma irradiation or ethylene oxide and provides more robust microbial clearance compared to UV exposure or ethanol disinfection^27^, demands the thermoformed material have a high glass transition temperature (T_g_). To thermoform a material with high T_g_, the tooling material used to create the mold must withstand high temperature for multiple cycles without cracking or exhibiting thermal deformation. Furthermore, many thermoplastics are hygroscopic, causing them to absorb moisture from the environment, which creates bubbles during thermoforming. This in turn compromises optical transparency, along with the ability to create micro-scale features.

An ideal material for thermoforming-assisted generation of cell culture devices would possess optical clarity, mechanical strength, low autofluorescence, biocompatibility, chemical inertness, support for additional cell adhesive coatings (e.g. extracellular matrix proteins like fibronectin, or protein cocktails Matrigel), and absence of harmful chemical leachates^28^. Polycarbonate (PC) emerges as a viable choice among thermoformable plastics, meeting these criteria (Table 1). PC has optical and mechanical properties similar to PS but can withstand autoclaving because of its high T_g_; biocompatibility of PC is demonstrated by its frequent use as a material of choice in making porous well plate inserts in cell culture (e.g. Corning Transwell products).

**Table 1.**
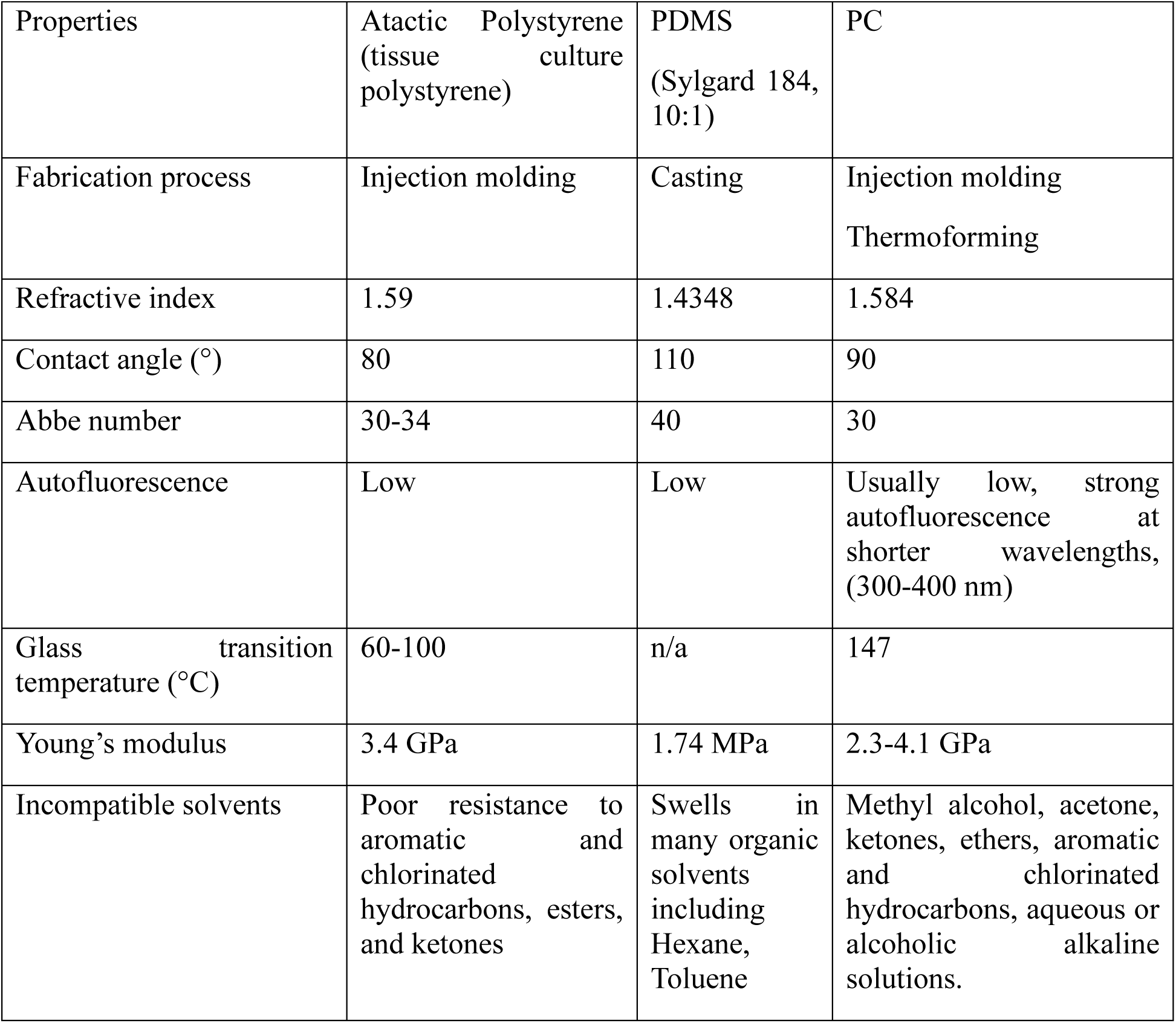
Mechanical and chemical properties of tissue culture polystyrene, PDMS and PC. ^32,33^

A key challenge in PC thermoforming lies in selecting appropriate tooling materials and fabrication conditions. Metal molds^29^ can withstand high temperature without thermal distortion or significant surface cracking. However, machining metal molds requires special skills, and expensive machinery, making it less suitable for a biologically focused research lab. Moreover, the need for frequent, iterative changes in the design of cell culture devices (**Fig. 1**), particularly during early prototyping stages, creates a need for alternative manufacturing methods that offer lower cost, more rapid design iteration.

Extrusion 3D printing (Fused Deposition Modeling, FDM) is one method to make mold rapidly^30^. However, because FDM requires heating the mold material to a flow-able state, materials used in FDM, like Polylactic acid, have low T_g_ (60-65°C) by design; they are thus not suitable for PC thermoforming. Even though other 3D printing techniques like stereolithography (SLA) and digital light processing (DLP) do not rely on the glass transition of the material for printing, most commonly available resin materials have T_g_ of ≤85°C. Recently, a specialized class of resins called high temperature deflection (HDT) resins has been developed for low run injection molding, automobile and dental applications^31^. HDT resins **(Table S1)** are capable of withstanding temperatures ranging from 180°C to 300°C^31^. As most of these resins are stable above the glass transition temperature of PC, these resins are a good choice for the tooling material for PC thermoforming.

In this study, we demonstrate the efficacy of combining DLP printing, high HDT resins, and desktop vacuum thermoforming for rapid fabrication of custom devices, culture platforms, and accessories tailored to various studies with iPSC and iPS-CM. We first show that optimizing preheating of PC sheets helps to circumvent bubbling during thermoforming, allowing creation of optically transparent devices with user-device micro-textures that we leverage for scalable formation of physiologically relevant 3D cardiobodies from iPSC. We anticipate this technology will support a variety of bioengineering and cell biology experiments.

## RESULTS

### High HDT tooling and preheating PC sheets helps to create transparent PC inserts with good dimensional accuracy

We first assessed the feasibility of PC thermoforming off molds made from several candidate tooling materials: aluminum, PLA, and high HDT resin). For these studies, we chose to replicate custom designed lids for a commercially available 70 mm petri dishes (Corning). Such lids find application in studies where external accessories must be placed close to the cell culture surface – for example, in our prior work, magnets were placed close to the surface of magnetically-responsive PDMS substrates to elicit dynamic changes in substrate elasticity^12^, while imaging through the bottom of the cell culture surface with an inverted microscope. This required a custom lid, as standard well plate lids create too large a gap between the magnet and the substrate. For this application to be feasible, the custom lid must fit precisely above the well surface, allowing gas exchange without excessive evaporation.

PLA molds failed to keep shape integrity during thermoforming, compromising the dimensional accuracy of the PC part. In contrast, machined aluminum and high HDT molds both withstood high temperature cycles without any visible thermal distortion, leading to production of nearly identical PC devices (**Fig. 2**). Subsequent studies confirmed that among available DLP resins, only high HDT resins could produce molds that could withstand many cycles of PC thermoforming without accumulating defects in the mold (**Fig. S1**). A Comparison of the mechanical and thermal properties of different DLP resins are listed in supplementary table 1.

Like many other thermoplastics, PC is hygroscopic ^34^ and absorbs moisture even with the relatively low humidity levels found in typical lab environments. At high temperature, these water molecules inside the PC expand, causing bubbles to form while thermoforming. We hypothesized that preheating the PC sheets could help to dry up the moisture prior to thermoforming, to minimize. Consistent with this hypothesis, PC sheets thermoformed after preheating did not exhibit bubbling, leading to improved optical transparency. Consistent with heat transfer scaling with the square of material thickness^35^, thin 0.03” PC sheets could be thermoformed without bubbling after 1.5hr preheating to 60°C, whereas thicker (0.04”) sheets required kept 5-6 hours of preheating. **(Fig. 3)**. PC sheets thickness less than 0.02” could be successfully thermoformed without bubbling without any preheating. This likely reflects the ability of these thinner sheets to heat rapidly enough during the thermoforming process to evaporate any surface moisture before the PC itself becomes pliable.

**Fig. 3.**
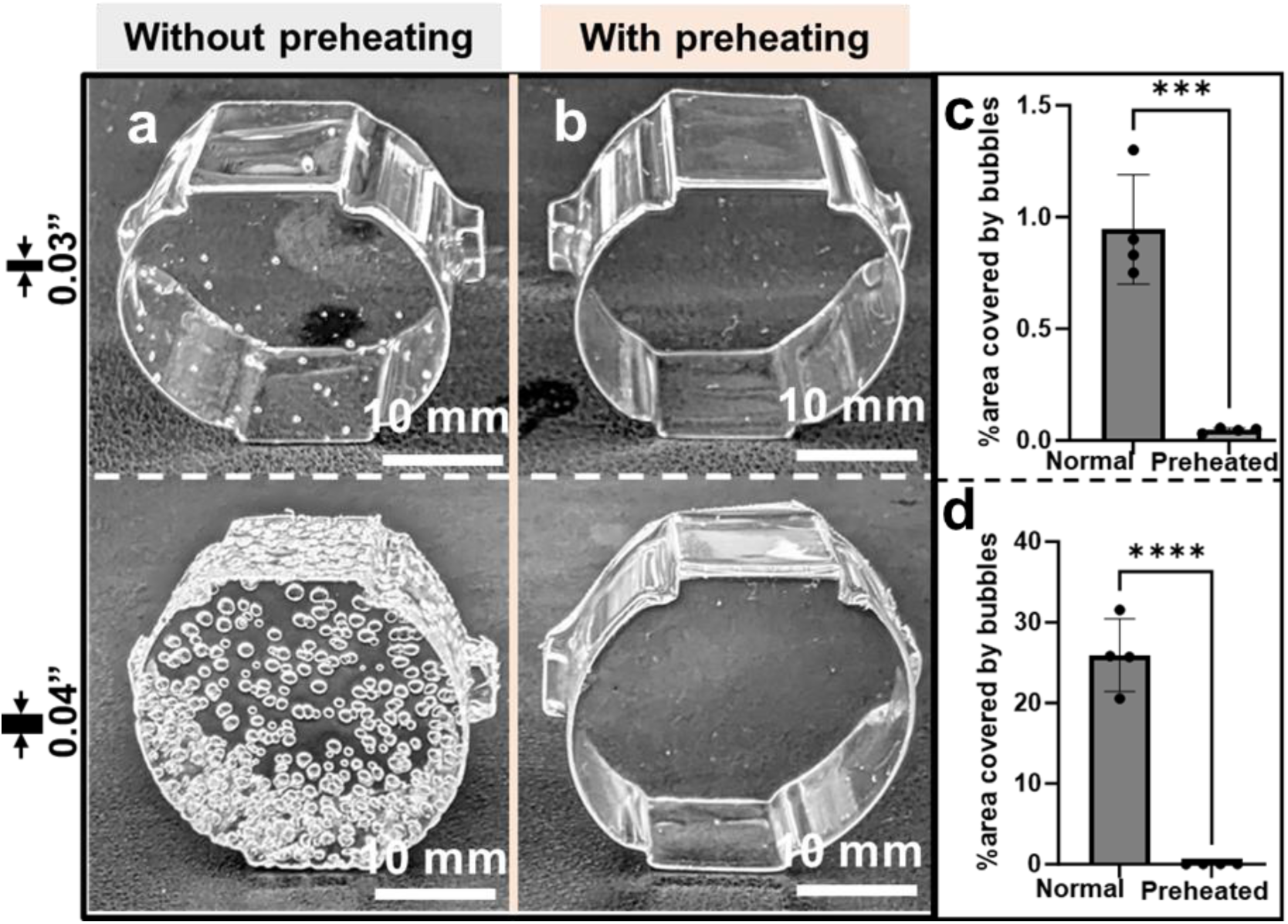
Preheating PC eliminates bubble formation during thermoforming. **A, B)** Representative images of PC inserts with different thickness that were thermoformed either A) without or B) with pre-heating for an optimized timeframe. **C, D)** Quantification of the percent area of the final thermoformed insert covered with bubbles when thermoforming was done either with or without preheating the PC. Each datapoint represents one thermoformed device. Error bars: *SD*, *** *p*<10^-3^.

### Thermoformed PC devices show good optical clarity, support fluorescence imaging, and do not absorb small hydrophobic molecules

One of the essential characteristics of plastic materials used in bioengineering studies is optical transparency to facilitate high-resolution imaging^36^. The emission and absorption spectra of well inserts made from PC or PDMS were similar to spectra of standard tissue culture polystyrene (PS; **Fig. 4a, b**), although PC outperformed PDMS, and PS had the highest optical transmittance. Absorption of hydrophobic molecules is a noted drawback of PDMS^37^. As expected, the hydrophobic probe molecule Rhodamine 6G absorbed into PDMS inserts (**Fig. 4c,d**). This led to a strong fluorescence emission peak at 560nm. In contrast, PC inserts did not absorb Rhodamine 6G, leading to negligible changes in optical transparency and fluorescence emission after applying this probe compound (**Fig. 4c, d,e**).

**Fig. 4.**
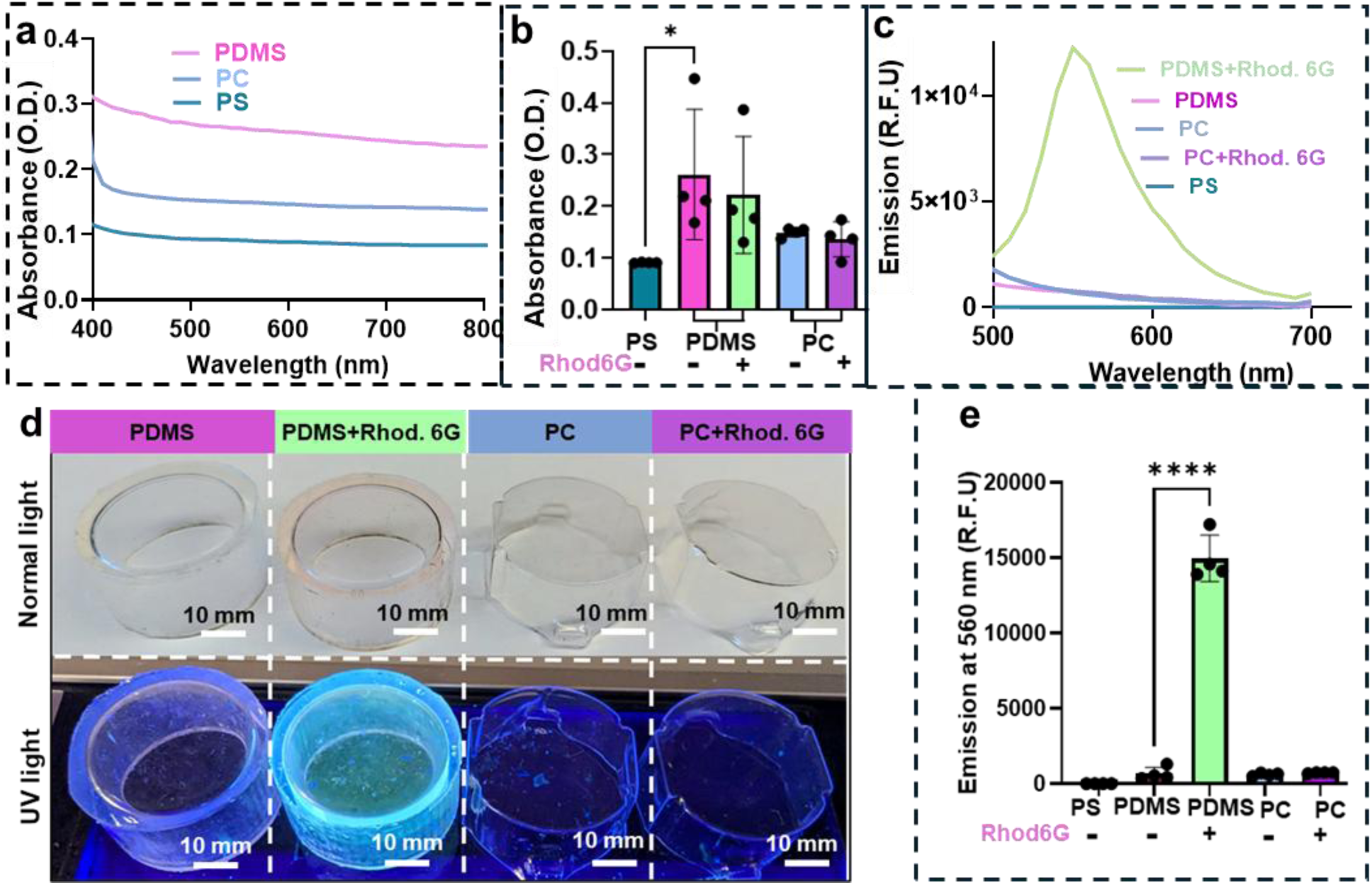
Thermoformed PC devices are optically transparent and resist absorption of hydrophobic molecules. **A)** Optical absorbance spectra of PDMS, PC and PS substrates of equivalent thickness (750 µm). **B,C)** Analysis of absorption of rhodamine G into the substrates through measurements of B) absorbance at 560nm (absorption peak of rhodamine G) or **C)** fluorescence emission spectrum after excitation at 490nm for PS, PDMS and PC either with or without pre-exposure to (1 mM) solution of rhodamine G. **D)** Representative images of molded PDMS devices, or thermoformed PC devices, either before or after exposure to 1mM rhodamine G solution. Devices were illuminated with UV light (365 nm). **E)** Quantification of peak fluorescence emission for rhodamine G (560nm) in PC and PDMS devices exposed to rhodamine G solution. Scale bars: 10 mm. Each data point represents one device. Error bars: *SD*. * *p* <0.05, **** *p* < 10^-4^, Holm–Šidák test following global ANOVA.

### C2C12 myoblasts and human induced pluripotent stem cells (iPSCs) exhibit good viability on thermoformed PC substrates

For PC to be used a cell culture substrate, it must facilitate robust cell adhesion and growth. Prior studies noted potential challenges in cell adhesion to PC substrates^38^. Consistent with these prior reports, we found that although C2C12 myoblasts cultured on PC substrate and tissue culture polystyrene were morphologically similar (**Fig. 5a**), overall cell attachment was lower on these uncoated PC substrates. This may reflect issues in adsorption of bioactive vitronectin, one of the main extracellular matrix proteins of cell culture serum that facilitates cell adhesion^39^, onto the PC. We also noted challenges in C2C12 adhesion to uncoated PDMS, consistent with prior reports that this material requires changes in surface chemistry to facilitate robust cell adhesion^40^. In contrast, C2C12 adhesion to PC substrates pre-coated with Geltrex, a commercial protein cocktail that mimics Matrigel (where the main extracellular matrix component is laminin)^41–43^ was indistinguishable from C2C12 adhesion to Geltrex-coated PS (**Fig. 5b**).

**Fig. 5.**
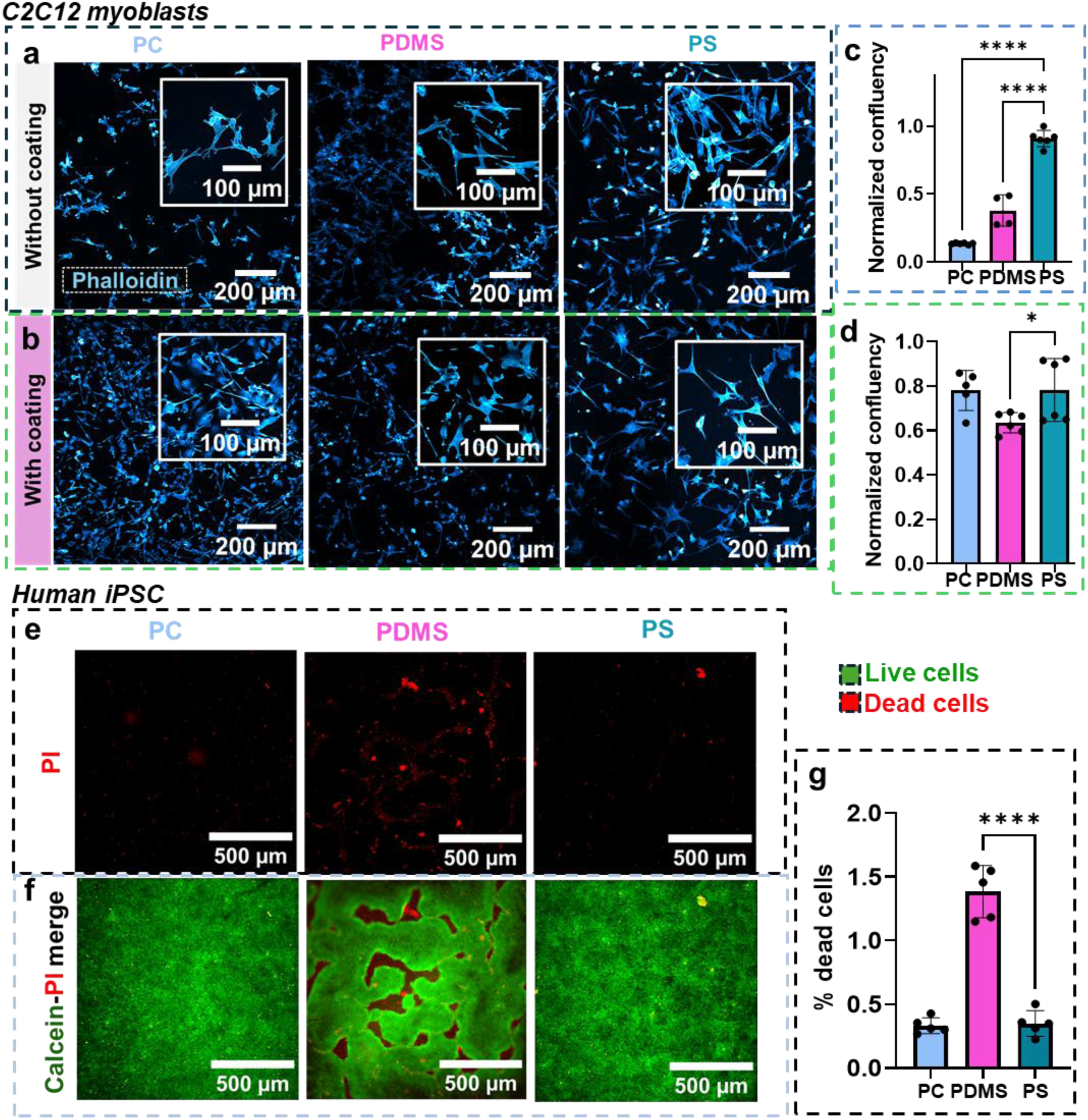
Extracellular matrix coated PC supports robust adhesion of myoblasts and induced pluripotent stem cells. **A, B)** Representative micrographs of C2C12 myoblasts when cultured on PC, PDMS and PS either A) without or B) with a coating of x ug/cm^2^ Geltrex. **C, D**) Quantification of overall cell adhesion (relative confluency, or area occupied by cells) of C2C12 on PS, PC or PDMS either C) without or D) with Geltrex coating. **E, F)** Representative E) calcein-AM (green) and propidium iodide (PI; red) co-staining and F) PI staining alone, for human iPSC cultured on Geltrex coated PS, PC or PDMS. **G)** Quantification of the area of dead (PI-positive cells). Error bars: *SD*. Each data point represents one ROI. Error bars: *SD*. * *p* <0.05, **** *p* < 10^-4^, Holm– Šidák test following global ANOVA.

**Fig. 6.**
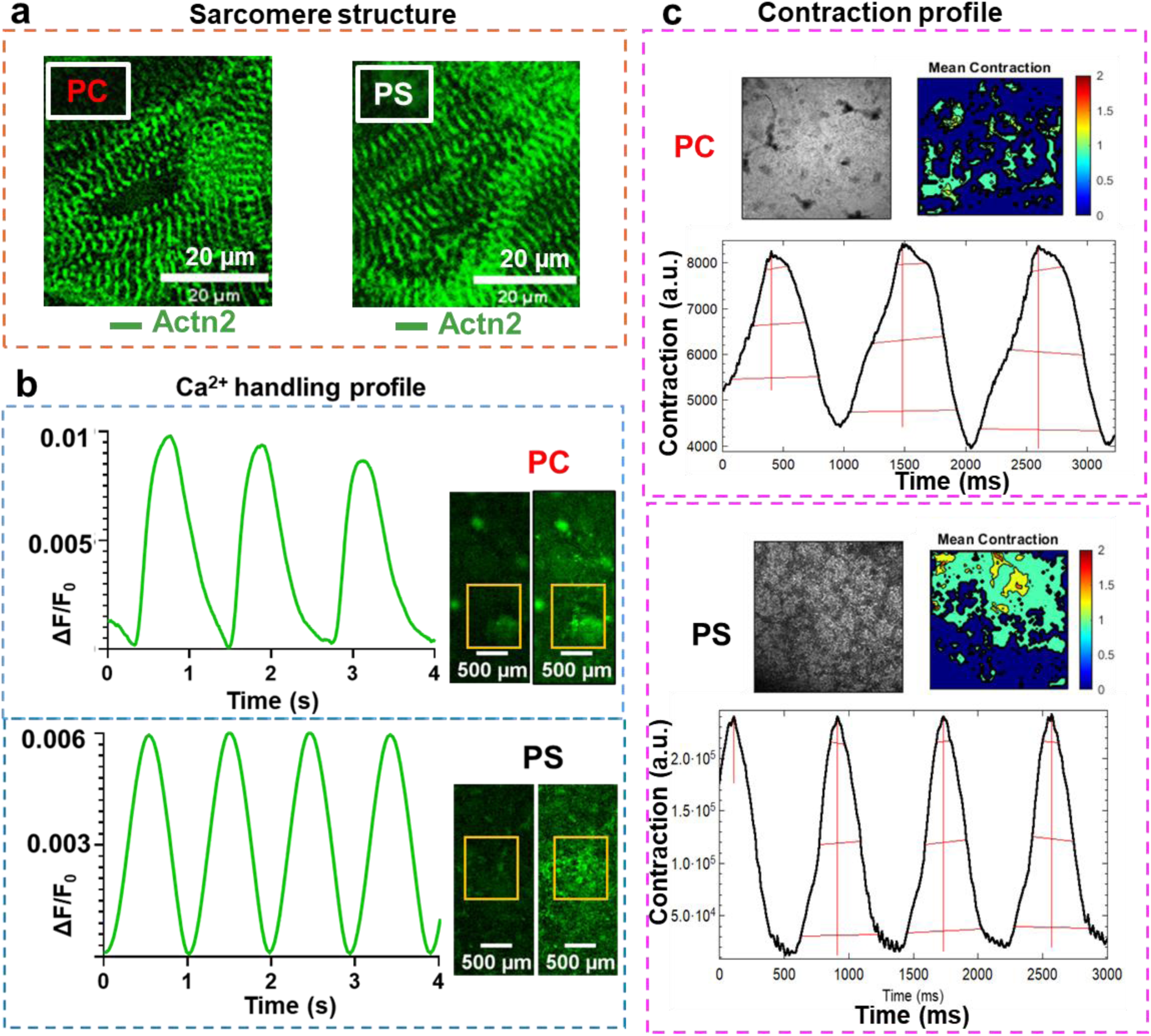
Geltrex-coated PC substrates support iPSC-cardiomyocyte differentiation. **A-C)** Representative A) micrographs of sarcomere organization, B) Ca^2+^ transient profiles and C) contractile motion profiles in iPS-CM sheets differentiated on (top) PC inserts or on (bottom) standard PS substrates. Insets in B) depict the GCaMP6f reporter in regions of the beating cardiomyocyte sheet during diastole (relaxed state, low cytoplasmic Ca^2+^) or systole (contracted state, high cytoplasmic Ca^2+^). See Supplementary video 1, 2.

While immortalized cells like C2C12 myoblasts are typically amenable to culture on a variety of different materials, pluripotent cells like iPSC typically have more stringent requirements in terms of culture substrate. For example, prior work shows that Matrigel coated tissue culture polystyrene, but not glass, supports robust human pluripotent stem cell attachment and growth^42^.

We thus tested the feasibility of using Geltrex coated PC for iPSC culture. Interestingly, we observed no significant changes in iPSC colony morphology or viability when these cells were cultured on Geltrex coated PC or PS (**Fig. 5b)**. In contrast, without chemical modifications, Geltrex-coated PDMS appeared to be slightly less suitable for supporting iPSC, with a small but statistically significant increase in observed cell death.

Aside from their ability to support C2C12 and iPSC adhesion, we also noted that PC inserts could withstand multiple cycles of sterilizing (bleaching, followed by washing and autoclaving), with no visible increase in cell death in iPSC cultured on PC substrates subjected to two cleaning and sterilization cycles. In contrast, after bleaching, PDMS failed to support cell growth even when the inserts were washed extensively before autoclaving (**Fig. S2**). While milder methods of disinfection may preserve PDMS, the ability to use a harsh sterilization technique like bleaching demonstrates the relative chemical resistance of PC.

### Thermoformed PC inserts support stem cell differentiation

iPSC differentiation to cardiomyocytes (iPS-CM) is a particularly demanding application for cell culture materials^44,45,46^ as cells undergo dramatic changes in morphology and adhesion to their culture substrate during this process. Moreover, as with any differentiation pathway, successful differentiation of iPSC down a cardiomyocyte lineage demonstrates the ability of the culture system to support these cells in their pluripotent state. We thus tested the ability of PC inserts to support this differentiation process. Surprisingly, we observed no gross changes in Ca^2+^ handling, beating patterns or sarcomere organization of iPS-CM differentiated atop Geltrex-coated PC versus standard PS substrates (**Fig, 6 a-c**).

### Thermoforming can create complex surface texturing in PC

3D printing can generate complex textures on the tooling material, and surface texture has marked effects on cell biology^47^. Thus, we next tested the feasibility of replicating complex textures on thermoformed PC. Using a simple dental-grade vacuum-forming machine (**Fig. 2**), the minimum achievable pattern dimension we achieved thermoformed PC was 250 μm (**Fig. 7a**). Above this lower limit, we successfully formed large areas of patterned PC substrates (**Fig. 8a-d**). iPSC cultured on patterned PC not only retained viability but grew in an ordered fashion (**Fig. 8a)**. We did note a substantially higher rate of cell death (propidium iodide positive cells; **Fig. 8b**) in the “ridges” formed by elevated features in the PC. We ascribe this to higher local cell density within the “valleys” between these ridges, as iPSC survival is strongly linked to local cell-cell contact^48^.

**Fig. 7.**
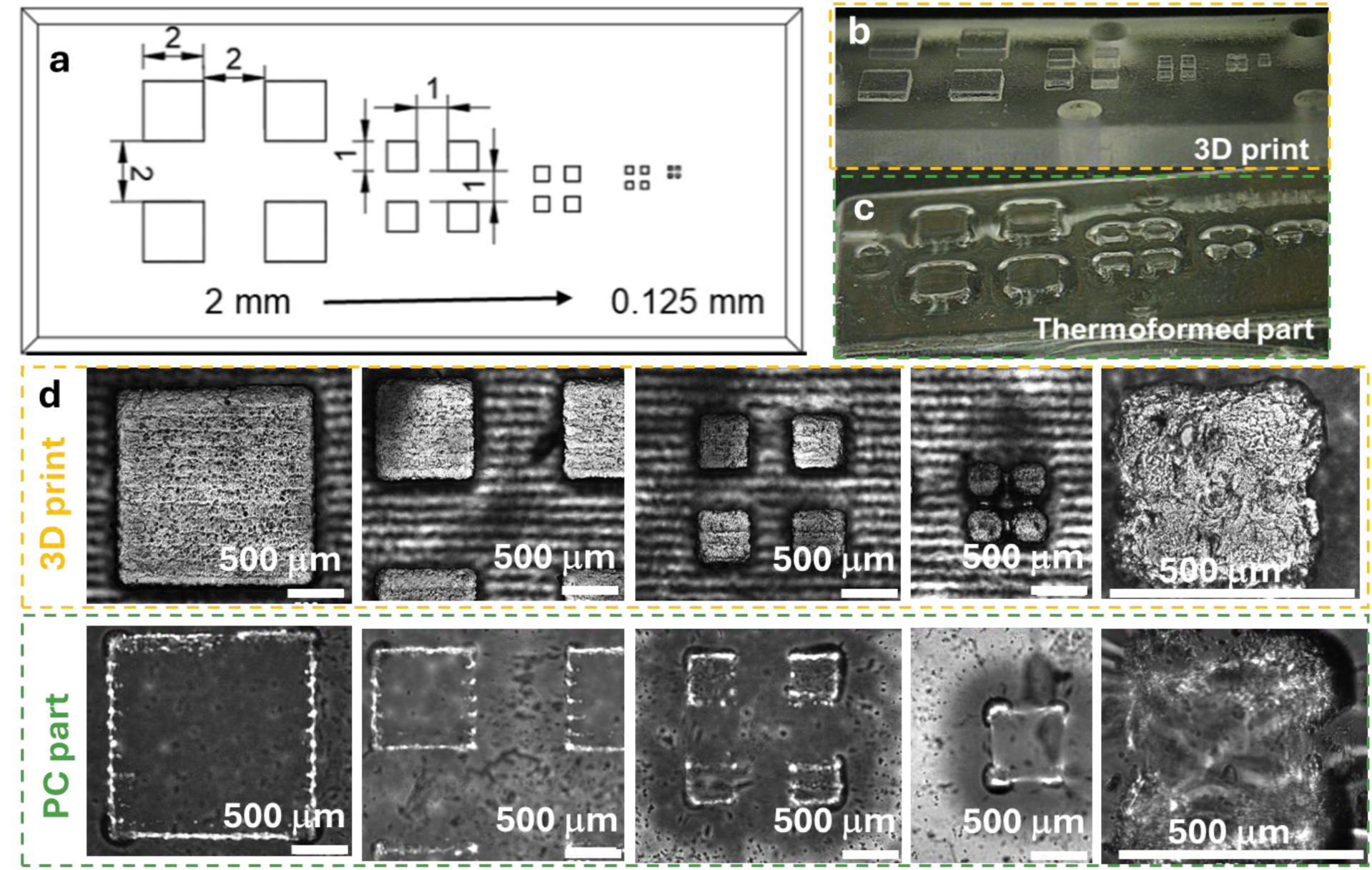
PC thermoforming can fabricate micropatterned arrays. **A)** rectangular array design for the fidelity test **B), C)** Photograph of B) 3D printed pattern, C) PC thermoformed part **D)** Micrographs show the quality of pattern replication

**Fig. 8.**
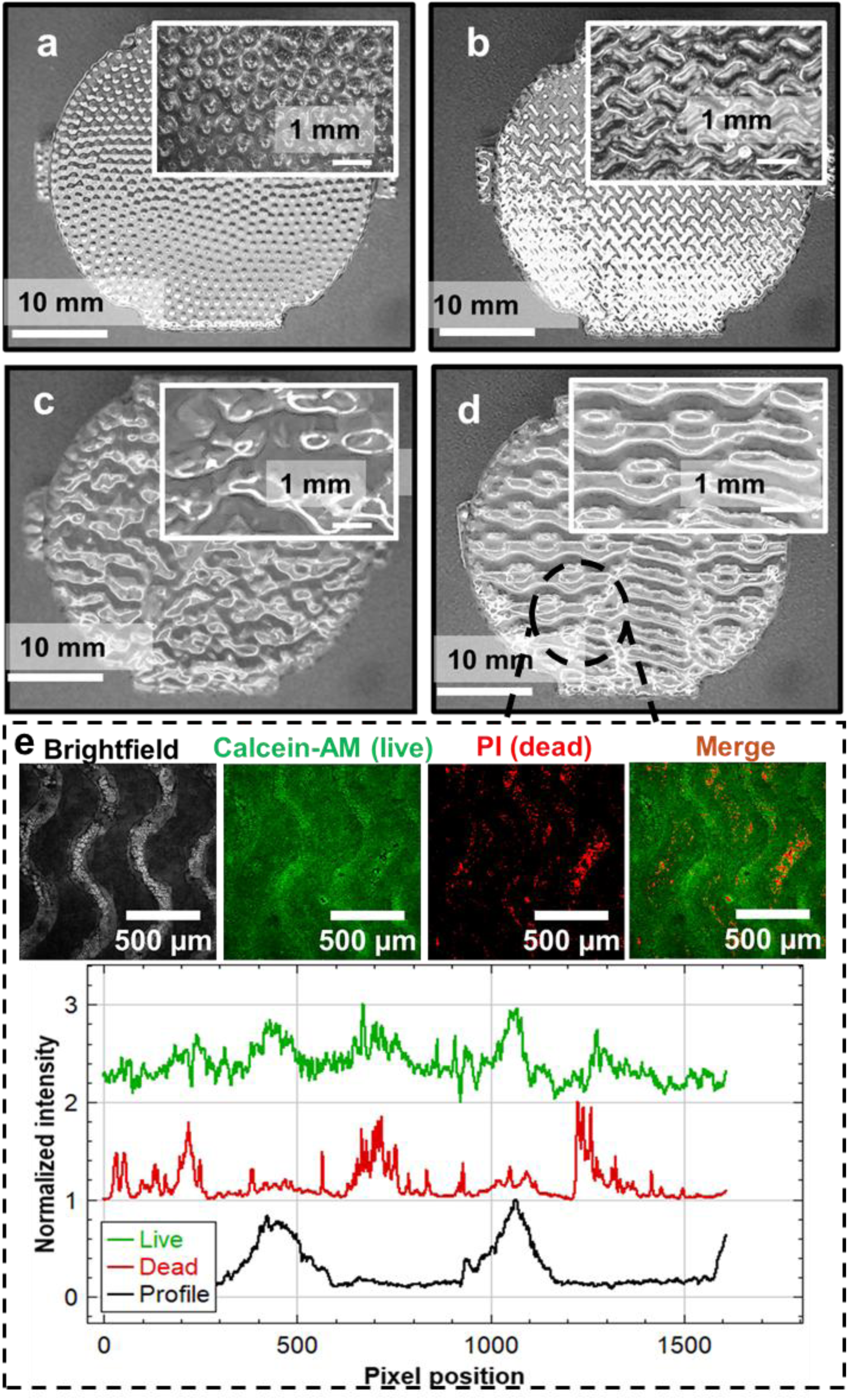
Thermoforming enables patterning cell culture surfaces **A)** bubble emboss pattern, **B)** checker plate pattern **C)** high roughness pattern **D)** wavy ridge pattern **E)** patterned inserts show difference in live/dead cell density distribution according to the surface topology. Scale bars: A-D: 10mm (insets: 1mm); E: 500μm.

### Thermoformed PC micro well inserts facilitate cell aggregation into spheroids

One of the main concerns of using iPSCs for modeling various diseases or conditions is about the lower level of maturity of iPSC-derived tissue cells compared to their postnatal counterparts^2^. 3D culture either during or after the differentiation process has emerged as a promising approach to improve iPS-CM maturity. Currently, there are limited options for the culture plates for forming spheroids^11,49,50^. Using PC patterned with pyramidal microwells, we successfully patterned C2C12 myoblasts (**Fig. 9a,b**) and iPS-CM **(Fig. 9c,d**) into spheroids. Aggregates formed within 3 days of seeding and could be cultured for at least 2 weeks.

**Fig. 9.**
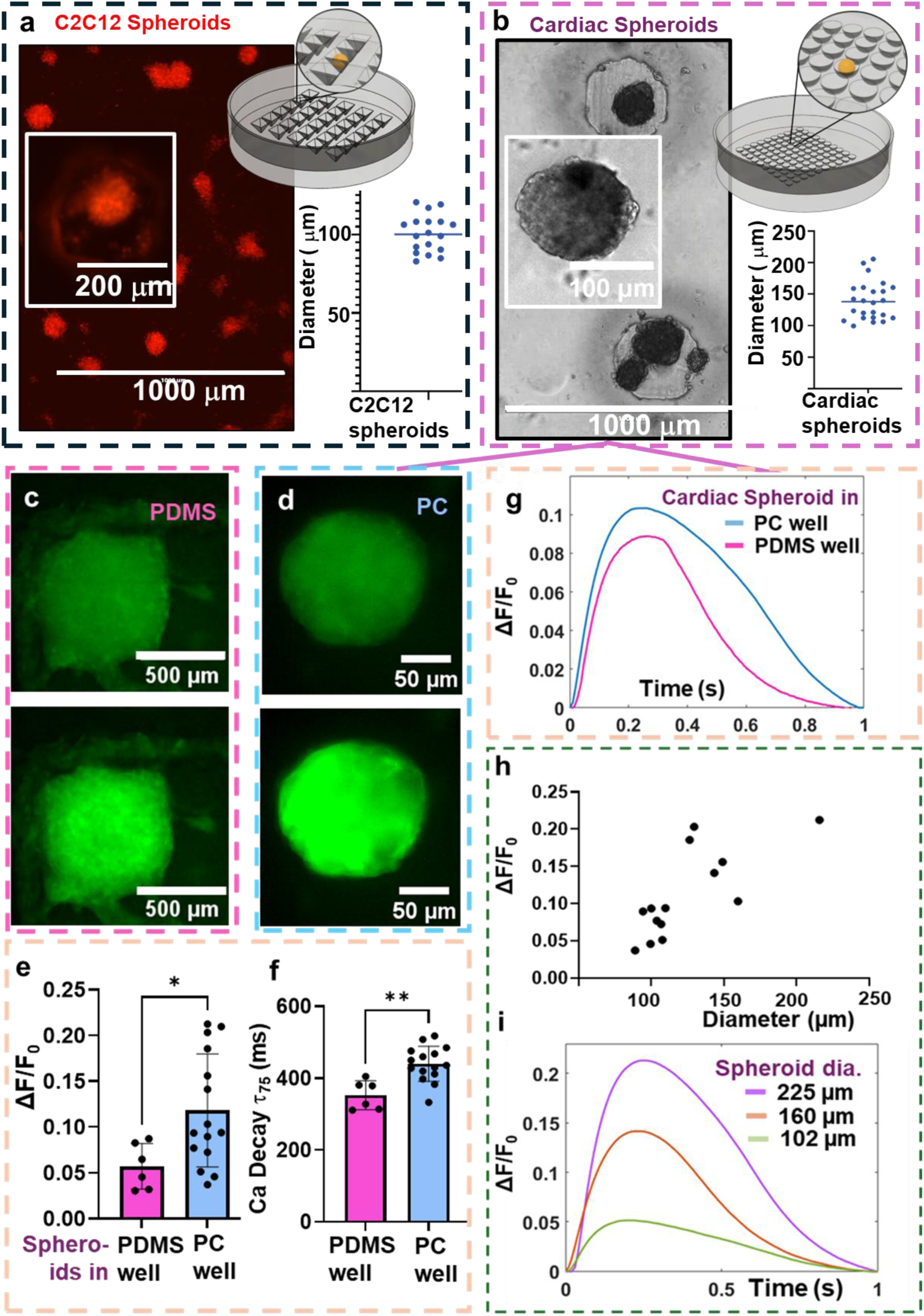
Leveraging PC micro-patterning to probe the relationship between cardiobody size and calcium handling dynamics. Micro patterned PC inserts facilitate spheroid formation A) C2C12 myoblast spheroids in a pyramid shaped microwell, B) cardiac spheroids in a hemispherical microwell C, D) representative images of spheroids formed in PC and PDMS wells during diastole (relaxed state, low cytoplasmic Ca^2+^) or systole (contracted state, high cytoplasmic Ca^2+^). E, F), G) show the Ca^2+^ transient analysis (background corrected Ca^2+^ intensity and time to reach 75% Ca^2+^ decay). H, I) demonstrate the difference in Ca^2+^ intensity with the change in cardiobody size. See supplementary video 3-6. Error bars: *SD*. * *p* <0.05, ** *p* <0.01, by unpaired t-test.

As our prior studies suggest a strong relationship between the geometry of engineered heart muscle and iPS-CM physiology, we addressed the impact of cardiobody geometry on calcium handling^46^. Compared to prior spheroid-shaped iPS-CM tissues formed in PDMS microwells ^46^, iPS-CM formed into cardiobodies in patterned PC wells exhibited greater baseline-corrected Ca^2+^ intake (ΔF/F_0_; **Fig. 9c,e,g**). The aggregates formed on the PC wells also exhibited a more spherical shape compared to those formed in PDMS microwells atop tissue culture plastic (**Fig 9c,d)**. We hypothesized that this stems from the larger size of cardiobodies formed in the PC microwells used in the present study. This hypothesis is consistent with the direct observation that Ca^2+^ intake increases, and the time to decay of individual Ca^2+^ transients becomes longer, as cardiobody size (diameter) increases (**Fig. 9d**).

## DISCUSSION

In this study, we demonstrate the feasibility of producing custom cell culture accessories from thermoformed PC that are transparent, mechanically and chemically stable, and which support cell culture and fluorescence imaging and even re-use after chemical sterilization, without heavy capital investment on machineries. Optimization of PC preheating time and temperature minimized bubbling down to a negligible level, ensuring optical transparency and mechanical stability, whilst allowing user-defined micro-patterning. This in turn allowed us to investigate a relationship between the geometry and Ca^2+^ handling of iPSC-derived cardiobodies. While a similar relationship could be studied in patterned PDMS, PC does not exhibit the same non-desirable absorption of small hydrophobic molecules.

As the custom cell culture accessories might be designed to support cell growth as a substrate or to “house” various subsystems inside the well keeping the environment sterile, it becomes critical to ensure that thermoformed PC parts are biocompatible. Our studies show that human iPSC, one of the more challenging cell types to maintain in culture, can be expanded and even differentiated on the PC inserts. Compared to iPS-CM differentiated on PS plates, PC inserts did not show any change in morphology or contractility (See supplementary Video).

DLP printing can create complex textures on the mold surface. As the study of the effect of substrate texture on cell behavior has a growing research interest, we attempted to fabricate PC inserts with different texture. The results show that the cells follow the textured patterns. In future studies, it will be of interest to combine PC’s abilities to support spatial patterning and differentiation to pattern development of organoids or differentiating monolayers^52,53^.

Calcium transient analysis of the cardiobodies shows significant increase in Ca2+ amplitude when cultured in PC microwells compared to PDMS wells. This may be related to changes in maturation, either of all cardiomyocytes within larger cardiobodies, or of the cardiomyocytes within specific regions of these larger cardiobodies. This may be related to gradients in nutrient availability and/or cell-synthesized extracellular matrix properties within the cardiobodies, and merits future studies.

Vacuum thermoforming has some inherent advantages and limitations in the context of cell culture device fabrication. In terms of advantages: besides the aforementioned lack of absorption of small hydrophobic molecules, PC also exhibited the ability to be sterilized by treatment with bleach, followed by washing and autoclaving, before re-use (**Fig. S2**). In some cases, it would be ideal to have the option of re-using cell culture accessories and/or substrates, particularly when these accessories are made from costly materials or have complex functions (e.g. if they incorporate devices for mechanical actuation^12^ or electrical pacing)^51^; this option is allowable with the present PC-thermoforming method.

In terms of disadvantages of the present approach and of the use of PC in particular: as PC has low flowability during injection molding, it is not recommended to reduce the wall thickness less than 1mm^54^. Contrary to that, thermoforming can achieve the wall thickness as low as 100μm which makes it suitable to make attachments that support high resolution imaging. However, replicating mold designs with large overhangs, with blind holes, with microscale texturing is difficult in thermoforming. Getting sharp corners is also challenging. Most of the designs may have to provide a draft angle of 1-2° for easy release of the part from the mold. Pressure thermoforming can address most of these drawbacks (except the need of draft angle).While the rigid mechanical properties of thermoformed PC are highly advantages for handling, specific applications that require substrates with mechanical properties tuned either to control cell biology or measure contractility may require softer materials, or coating the PC materials with a thin layer of soft materials such as hydrogel or PDMS.

Finally, we note that PC is susceptible to photodegradation during extended UV irradiation especially in the presence of oxygen^55^. Therefore, it is better to avoid UV sterilization of the PC attachments when they used for cell culture applications. This drawback is overcome by the ability to apply autoclaving, which is more robust at eliminating microbes compared to UV.

## CONCLUSION

We suggested a method for rapid fabrication of custom cell culture/tissue culture accessories by combining DLP printing, high HDT resins and thermoforming PC sheets. As a 0.03” PC sheet takes 20 seconds to soften and it takes less than one minute to fabricate a part, and the mold can be reused for at least 1000 cycles without any visible thermal damage, this creates a low-cost approach for creating bespoke cell culture/tissue engineering accessories with medium throughput. We further demonstrated that thermoformed PC devices are capable of facilitating cell growth and fluorescent imaging. As 3D cell culture and co-culture are proving critical in many in vitro disease modeling studies ^2,56^ researchers can use the simple approach presented here to design, fabricate and test new customized devices for their need.

We expect the flexibility of the PC thermoforming process will help researchers to iterate through device designs as an intermediate step before going for mass production approaches such as injection molding. In summary, this approach offers a rapid, cost-effective, flexible pathway for researchers to develop custom solutions for their tailored experiments where commercial products fall short.

## MATERIALS AND METHODS

### CAD and 3D printing

All the designs are made with Autodesk Fusion 360 and sliced with *CHITUBOX Basic* (Version 2.0). We used Phrozen Mighty 8K DLP printer and Siraya Tech Sculpt high HDT resin to print the thermoforming molds. Other DLP resins including Conjure rigid resin, Phrozen clear 8k resin, Phrozen gray 8k resin, TH47 tough resin are used to compare the performance of HDT resins in multiple cycles of thermoforming.

### Thermoforming

We used Plastvac P7 Dental Vacuum Forming Machine (Bioart, Brazil) for thermoforming PC parts. We used RowTec® Polycarbonate Film of thickness (0.03”, 0.04”). Thickness less than 0.03” was difficult to handle and not watertight due to micro cracks during thermoforming. Thickness greater than 0.04” was difficult to remove from the mold after thermoforming.

### iPSC-cardiomyocyte differentiation

All iPSC experiments were conducted with Wild Type C (WTC) human hiPSC line which express single copy of GCaMP6f (Coriell Repository # GM25256). These cells undergo at least 3 passages in Essential 8 medium before starting the differentiation protocol. On all the conditions (PC, PDMS, and PS), culture surface was coated with Geltrex™ (Thermo Fisher Scientific, Waltham, MA) diluted in KnockOut™ DMEM (Thermo Fisher Scientific) at a ratio of 1: 100 to ensure cell attachment. During first 4 days of differentiation, the cells were cultured in RPMI 1640 (Thermo Fisher Scientific) with B27 supplement without insulin and with 150 μg/mL L-Ascorbic acid (Sigma, Millipore Sigma, St. Louis, MO). Intermittently, additional reagents were supplied to guide them to cardiac lineage. After ensuring 75-80% confluency, the cells were treated with 6 μM CHIR99021 (Biogems) for GSK-3β inhibition. Wnt signaling inhibitor (5 μM IWP-2, Biogems). was supplied after 48hrs. On day 4, IWP-2 was removed, and the medium contained only R− and L-ascorbic acid. Beginning on day 6, the cells were maintained in RPMI 1640 supplemented with B-27 containing insulin (R+), with media refreshed on days 6, 8, and 11, and then every three days. Beating activity in the cell layer typically began to emerge between days 6 and 10.

For making cardiobodies, the cells are further replated and purified. On day 15 of differentiation, contracting cardiomyocyte layers were enzymatically dissociated using dispase and 1× TrypLE^TM^ Express (1X) for 15 minutes each. Cells were carefully pipetted in knockout DMEM containing 20% fetal bovine serum (Gibco) to achieve a single-cell suspension. The cells were then resuspended in RPMI 1640 medium supplemented with B27 (with insulin) and 10 μM Y27632 and transferred to Matrigel-coated plates. Beating activity typically reappeared within 24 hours. On day 16, the medium was refreshed with standard RPMI/B27. For metabolic purification, between days 20 and 24, cells were cultured in glucose-free medium supplemented with 4 mM sodium L-lactate (Sigma), promoting selective survival of cardiomyocytes. Cultures were then transitioned back to RPMI/B27 over the following 10 days, gradually increasing the proportion of RPMI/B27 to 100% over the final 3 days. The cells are used for cardiobodies within 3days after transition.

### C2C12 immortalized mouse myoblast culture

C2C12 myoblasts were seeded onto various surfaces at a density of 5,000 cells per cm² using Dulbecco’s Modified Eagle Medium (DMEM; Thermo Fisher Scientific) supplemented with 1% fetal bovine serum (FBS; Thermo Fisher Scientific).

### Cell viability

Cell viability was evaluated using Calcein-AM and Propidium Iodide (PI) staining (Live/Dead Imaging Kit, Thermo Scientific). Cells were incubated with 2 μM Calcein-AM and 4 μM PI in PBS at 37 °C for 30 minutes, protected from light. After washing with PBS, fluorescence imaging was performed using a fluorescence microscope. Live cells fluoresced green (Calcein-AM), and dead cells fluoresced red (PI). Quantification was done using ImageJ.

### Cardiobody formation

Purified cardiomyocytes are dissociated using TrypLE^TM^ Select 10X and replated on to the special PC inserts with RPMI 1640 medium supplemented with B27 (with insulin), 10% FBS and 10 μM Y27632. The inserts are centrifuged at 100 rpf for 4 minutes to allow the cells to aggregate to single microwells. The inserts are carefully transferred to the incubator and kept for 48hrs. The media changed carefully to RPMI 1640 medium supplemented with B27 (with insulin) after 48 hrs. Videos are captured after culturing the cardiobodies for 5 days.

### Calcium transient measurement

Intracellular calcium activity was monitored using the GCaMP6f calcium indicator. Time-lapse fluorescence imaging was performed at a frame rate of 100 fps to capture calcium transients. A custom MATLAB script^57^ was used to isolate individual calcium waveforms and quantify key parameters, including spontaneous beating frequency, signal amplitude, rise time, and decay kinetics.

### Contraction analysis

The contraction maps were generated using MotionGUI^58^ software and the contraction profile graphs were plotted using MUSCLEMOTION^59^.

### Immunostaining

Cardiomyocyte monolayers were fixed with 4% paraformaldehyde (PFA) for 2 hours at room temperature, followed by PBS washes. Cells were permeabilized using 0.1% Triton X-100 for 10 minutes, then blocked for 45 minutes in a solution containing 3% goat serum and 3% BSA in 0.1% Triton X-100. Samples were incubated with a primary antibody against sarcomeric α-actinin (ACTN2) and subsequently imaged using a confocal microscope (Olympus Fluoview FV1200, Tokyo, Japan).

### Statistical analysis

All data are presented as mean ± standard deviation (SD). For comparisons between two groups, statistical significance was assessed using unpaired two-tailed Student’s t-tests. For comparisons involving more than two groups, one-way ANOVA followed by Holm–Šidák post hoc tests was used to correct for multiple comparisons. A minimum of four biological replicates (independent inserts) were used for each condition. Statistical significance was defined as p < 0.05.

## Acknowledgments

We thank Michael Vahey for providing access to the confocal microscope. We are also grateful to lab members Hsin Yi Chou, Huanzhu Jiang, Yasaman Kargar Gaz Kooh, Ghiska Ramahdita, and Austin Kellogg for their valuable feedback during testing of the process for their specific applications. We also thank Mohammad Hashemi for his insightful comments. Finally, we thank Daniel Simmons for his help with generating and analyzing cardiobody data.

## Author contributions

Conceptualization: GM

Methodology: GM

Investigation: GM

Visualization: GM

Writing—original draft: GM, NH

Writing—review & editing: GM, NH

Supervision and Funding: NH

## Data and materials availability

All data are available in the main text or the supplementary materials.

## Ethics Approval

Ethics approval not required.

