## Supplementary Material text for "Rapid thermoforming of polycarbonate cell culture accessories from 3D printed molds"

### Supplementary Materials

| Table of contents: Supplementary materials | | | |
|---|---|---|---|
| **Label** | | **Type** | **Caption** |
| **Supplementary document (PDF)** | Supplementary Table S1 | Table | *Mechanical and thermal properties of mold materials* |
| | Supplementary Figure 1 (Fig. S1) | Image | *Extended analysis of the impact of tooling material composition on thermal and resistance.* |
| | Supplementary Figure 2 (Fig. S2) | Image | *PC inserts withstand but PDMS inserts fail the reusability test* |
| **Supplementary videos (GIF/AVI)** | Supplementary Video 1 | Video | *Differentiated iPS-CM on Tissue culture PS substrate* |
| | Supplementary Video 2 | Video | *Differentiated iPS-CM on PC substrate* |
| | Supplementary Video 3 | Video | *Brightfield video of cardiosphere formed in PC well* |
| | Supplementary Video 4 | Video | *Fluorescent video showing $Ca^{2+}$ handling in cardiosphere formed in PC well* |
| | Supplementary Video 5 | Video | *Brightfield video of cardiosphere formed in PDMS well* |
| | Supplementary Video 6 | Video | *Fluorescent video showing $Ca^{2+}$ handling in cardiosphere formed in PDMS well* |

| Table S1. Mechanical and thermal properties of mold materials[31,60–62] | | | | | | | |
|---|---|---|---|---|---|---|---|
| Property | Conjure Rigid Gray | Phrozen Aqua Clear | Phrozen Aqua Gray | TH72 | Siraya Sculpt High HDT | PLA (Polylactic Acid) | Aluminum |
| Type | DLP Resin | DLP Resin | DLP Resin | DLP Resin | DLP Resin | Thermoplastic | Metal |
| Ultimate tensile Strength (MPa) | 39.13 | 17.6 | 17.6 | 26.23 | 35 | 40 | 310 |
| Heat Deflection Temperature (°C) | 70 | 70 | 70 | 80 | 180 | 60 | 500+ |
| Shore Hardness | 85–90D | 70D | 70D | 75D | 90D | Not applicable | Not applicable |
| Transparency | Opaque | Clear | Opaque | Opaque | Clear | Opaque | Opaque (metallic) |

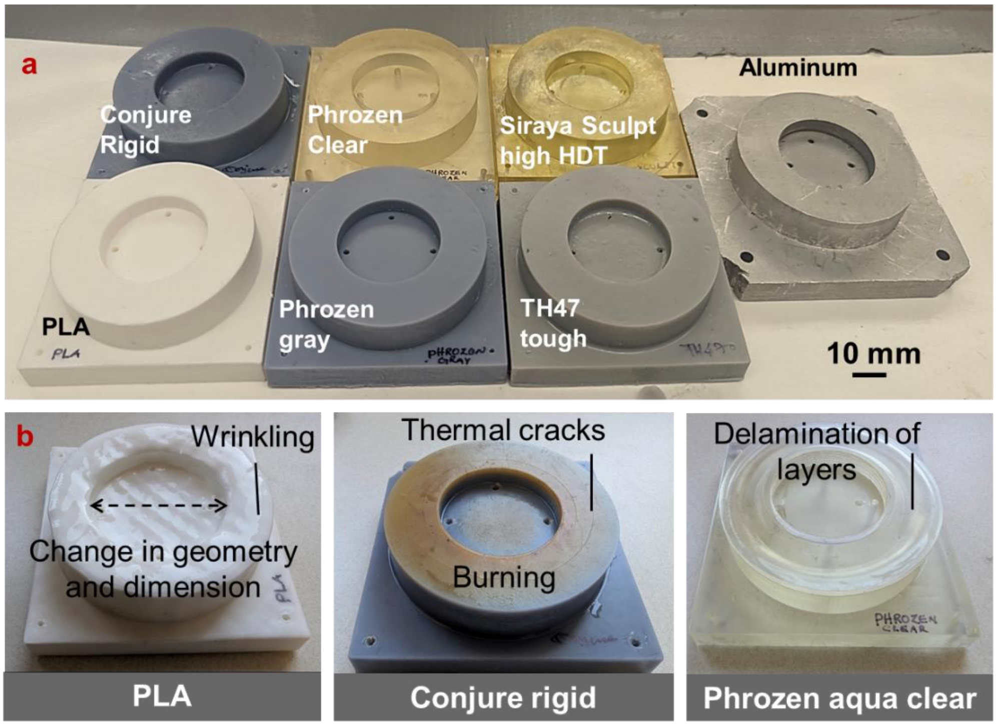


**Fig. S1. Extended analysis of the impact of tooling material composition on thermal and resistance.** a) Shows molds fabricated with different DLP resins, PLA and aluminum b) common thermal defects on the mold after being exposed to high temperature. Scale bar: 10mm.

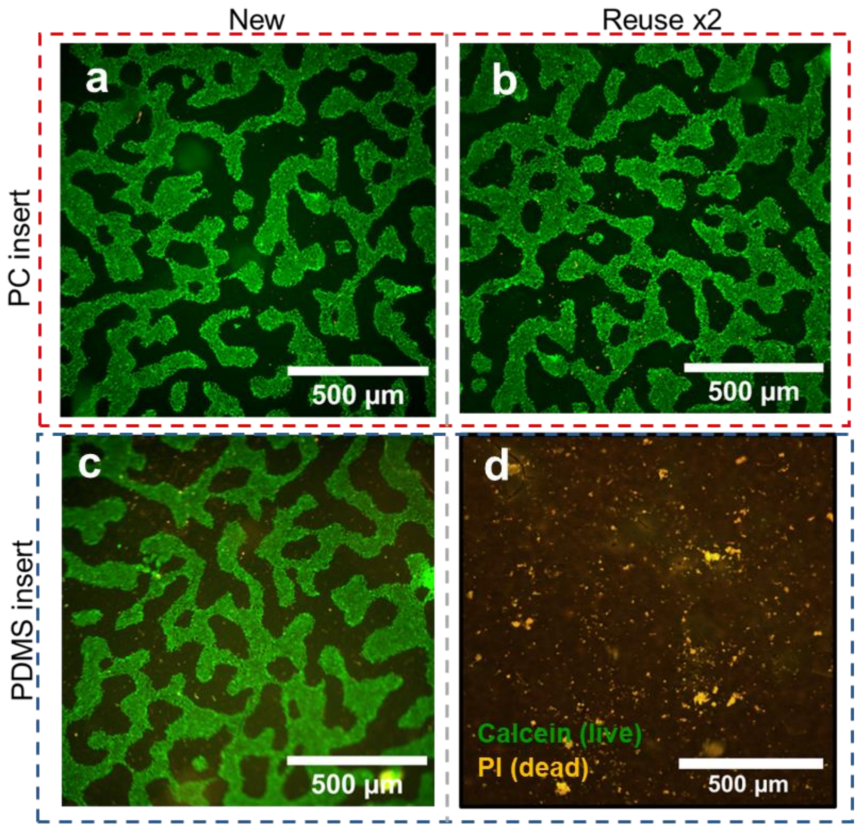


**Fig. S2**. **PC inserts withstand but PDMS inserts fail the reusability test** Representative calcein-AM (green) and propidium iodide (PI; red) co-staining show human iPSC cells grow similarly in **A)** fresh PC insert and **B)** reused PC insert, **C)** fresh PDMS insert, **D)** but limited cell growth in reused PDMS insert. Scale bars: 500µm.
